# Aspartyl Protease 5 matures virulence factors found at the host-parasite interface in *Toxoplasma gondii*

**DOI:** 10.1101/271676

**Authors:** Michael J Coffey, Laura F Dagley, Eugene A Kapp, Giuseppe Infusini, Justin A Boddey, Andrew I Webb, Christopher J Tonkin

## Abstract

*Toxoplasma gondii* infects approximately 30% of the world’s population, causing disease primarily during pregnancy and in individuals with weakened immune systems. *Toxoplasma* secretes and exports effector proteins that modulate the host during infection and several of these proteins are processed by the Golgi-associated Aspartyl Protease 5 (ASP5). Here, we identify ASP5 substrates by selectively enriching N-terminally-derived peptides from wildtype and Δ*asp5* parasites. We reveal over two thousand unique *Toxoplasma* N-terminal peptides, mapping to both natural N-termini and protease cleavage sites. Several of these peptides mapped directly downstream of the characterised ASP5-cleavage site, arginine-arginine-leucine (RRL). We validate candidates as true ASP5 substrates, revealing they are not processed in parasites lacking ASP5, nor in wild type parasites following mutation of the motif from RRL⟶ARL. All new ASP5 substrates are dense granule proteins, and interestingly none appear to be exported, thus differing from the analogous system in related *Plasmodium* spp., instead revealing that the majority of substrates reside within the parasitophorous vacuole (PV), and its membrane (the PVM), including two kinases and one phosphatase. Furthermore, we show that several of these ASP5-substrates are virulence factors, with their removal leading to attenuation in a mouse model, suggesting that phosphorylation at the host-parasite interface is important for virulence. Collectively, these data constitute the first in-depth analyses of the total list of ASP5 substrates, and shed new light on the role of ASP5 as a maturase of dense granule proteins during the *Toxoplasma* lytic cycle.

## Importance

*Toxoplasma gondii* is one of the most successful human parasites. Central to its success is the arsenal of virulence proteins introduced into the infected host cell. Several of these require direct maturation by the aspartyl protease ASP5, and all require ASP5 for translocation into the host cell, yet the true number of ASP5 substrates is currently unknown. Here we selectively enrich N-terminally-derived peptides using Terminal Amine Isotopic labelling of Substrates (TAILS) and quantitative proteomics to reveal novel ASP5 substrates. We identify, using two different enrichment techniques, new ASP5 substrates and their specific cleavage sites. ASP5 substrates include two kinases and one phosphatase that reside at the host-parasite interface, which are important for infection.

## Introduction

Apicomplexan parasites are the causative agents of many diseases of important medical and agricultural significance. As obligate intracellular pathogens, these parasites must invade and then survive within the infected host cell whilst obtaining nutrients in order to replicate. *Toxoplasma gondii* is the most widespread and successful of all apicomplexan parasites, and resides in nucleated cells of nearly all warm-blooded organisms, including birds and mammals. Initial infection in immunocompetent humans is generally mild, however some highly virulent South American strains of *Toxoplasma* exist and cause progressive blindness in otherwise healthy individuals (1). Further, reactivation of latent infection within immunocompromised populations, such as AIDS and immunotherapy patients, can cause severe disease and death (2).

Central to the success of *Toxoplasma* is its ability to modulate the host response, a process achieved through the secretion and export of an arsenal of effector proteins. Subversion of the host enables the parasites to survive and proliferate within the parasitophorous vacuole (PV), a structure that delimits *Toxoplasma* from the hostile intracellular environment. At the molecular level, *Toxoplasma* achieves this fine balance of immune modulation by the orchestrated secretion of effector proteins using two separate protein trafficking pathways. Proteins from the rhoptry organelles (termed ROPs) are introduced during the invasion process where they simultaneously promote a pro-inflammatory microenvironment to protect the host from excessive parasite growth, whilst also ensuring *Toxoplasma* is not cleared by the mounting immune response (3–11).

More recently, it has been shown that another protein export pathway exists. Here, dense granule proteins (GRAs) are secreted post-invasion and can modulate the host either by localising to the PV membrane (PVM) or by translocating across this barrier and residing within the host cell. GRA15, for example, localises to the host cytosolic side of the PVM where it activates the host NFĸB pathway to induce a protective immune response against the parasite (12), whilst the transmembrane protein MAF1 recruits host mitochondria to the PVM (13).

Another group of dense granule proteins are translocated across the PVM and into the host cell. GRA16 traffics to the host cell nucleus where it interacts with the phosphatase PP2A and the ubiquitin protease HAUSP, potentially interfering with the cell cycle to avoid premature immune detection (14, 15). GRA24, on the other hand, induces a protective response through the prolonged activation of the MAPK protein p38α leading to controlled replication within the gut of infected mice (15, 16). One of the major subversion mechanisms of *Toxoplasma* is the loss of the infected host cell’s ability to mount an IFNγ response. This response is circumvented by the dense granule protein IST (inhibitor of STAT1-dependent transcription). IST directly binds to activated STAT1 in the host nucleus and recruits a chromatin remodelling complex to block the transcription of STAT1-dependent promoters (17, 18). GRA28 was discovered in a screen by tagging a dense granule protein with the promiscuous biotin ligase BirA, although its role in the host nucleus has not yet been elucidated (19).

The precise mechanism of how *Toxoplasma* proteins translocate across the PVM is currently unclear; however, the export of GRA16, GRA24 and IST is dependent on the Golgi-resident aspartyl protease ASP5, which is known to directly process the exported protein GRA16, and likely IST. Despite not proteolytically maturing GRA24 (17, 20–22), export of this effector still requires the activity of ASP5. ASP5 cleaves its substrates at a defined motif termed the TEXEL (*<u>T</u>oxoplasma* <u>Ex</u>port <u>El</u>ement) a conserved motif consisting of RRLxx, so named as it is homologous to the cleavage of the *Plasmodium* export element (PEXEL, commonly RxLxE/Q/D) by the ER-resident protease plasmepsin V (PMV) (20, 23–27). Within malaria parasites, the PEXEL appears to only occur near the N-terminus of the protein and its cleavage by PMV somehow licenses proteins for export across the PVM via the *Plasmodium* translocon of exported proteins (PTEX) (24, 28–36).

Despite similarities, there are also several key differences between these pathways in *Toxoplasma* and *Plasmodium*. ASP5 resides within the Golgi, where it cleaves the TEXEL that can be found at the N- or C-terminus of the substrate. Further, while all reported PEXEL proteins are exported, it appears only a subset of ASP5 substrates are translocated into the host cell. We recently reported that ASP5 processes the PVM protein MYR1, a likely component of the *Toxoplasma* translocon (20, 37). Further, previous work by us and others demonstrated that ASP5-dependent effectors substantially alter the transcriptional profile of the host cell. Due to its central role in host cell subversion, parasites lacking ASP5 exhibit decreased virulence in mice, even in the normally lethal RH strain (20, 22).

Despite the importance of ASP5, only a handful of ASP5 substrates have been identified, and the true number of substrates remains unknown. Indeed, identification of new ASP5 substrates could identify new effector proteins and virulence factors driving the persistence of *Toxoplasma*. To identify new ASP5-dependent effector proteins, we have utilised a quantitative proteomic pipeline in combination with the selective enrichment of N-terminally derived peptides. These methods included Terminal Amine Isotopic Labelling of Substrates (TAILS) (38) and Hydrophobic Tagging-Assisted N-Terminal Enrichment (HYTANE) (39), which enabled us to compare differences in the N-terminome between wildtype (WT) and Δ*asp5* tachyzoites. Enrichment of N-terminal peptides by TAILS and HYTANE has enabled the identification of protease cleavage sites dependent on ASP5.

Moreover, we validated the N-terminal enrichment data and report that ASP5 matures several new dense granule proteins that appear to be localised within the confines of the parasitophorous vacuole. We present LCAT (40) and a new dense granule protein, GRA43, as ASP5 substrates that are processed close to the C-terminal end of the polypeptide. Further, we validate WNG1 and WNG2, formerly ROP35 and ROP34, as two new dense granule protein kinases processed by ASP5. This study greatly increases the number of known ASP5 substrates, including several involved *Toxoplasma* virulence, demonstrating the important role protein maturation plays in pathogenesis.

## Results

### Identification of ASP5 substrates by N-terminal enrichment

In our current model, we propose that *Toxoplasma* effectors enter the ER via an N-terminal signal peptide that is subsequently cleaved by signal peptidase. Proteolytic cleavage can result in acetylation (or not) of the new N-termini (Figure 1A). Upon reaching the Golgi apparatus, ASP5 matures substrates at the TEXEL (RRLxx), prior to transport across the parasite plasma membrane (PPM) (Figure 1A). Within the PV space, some substrates are inserted into the PVM, while others are exported into the host cell. Parasites that lack ASP5 are unable to mature these substrates within the Golgi, resulting in different N-termini, commonly represented as the signal peptide cleavage site (Figure 1A).

**Figure 1:**
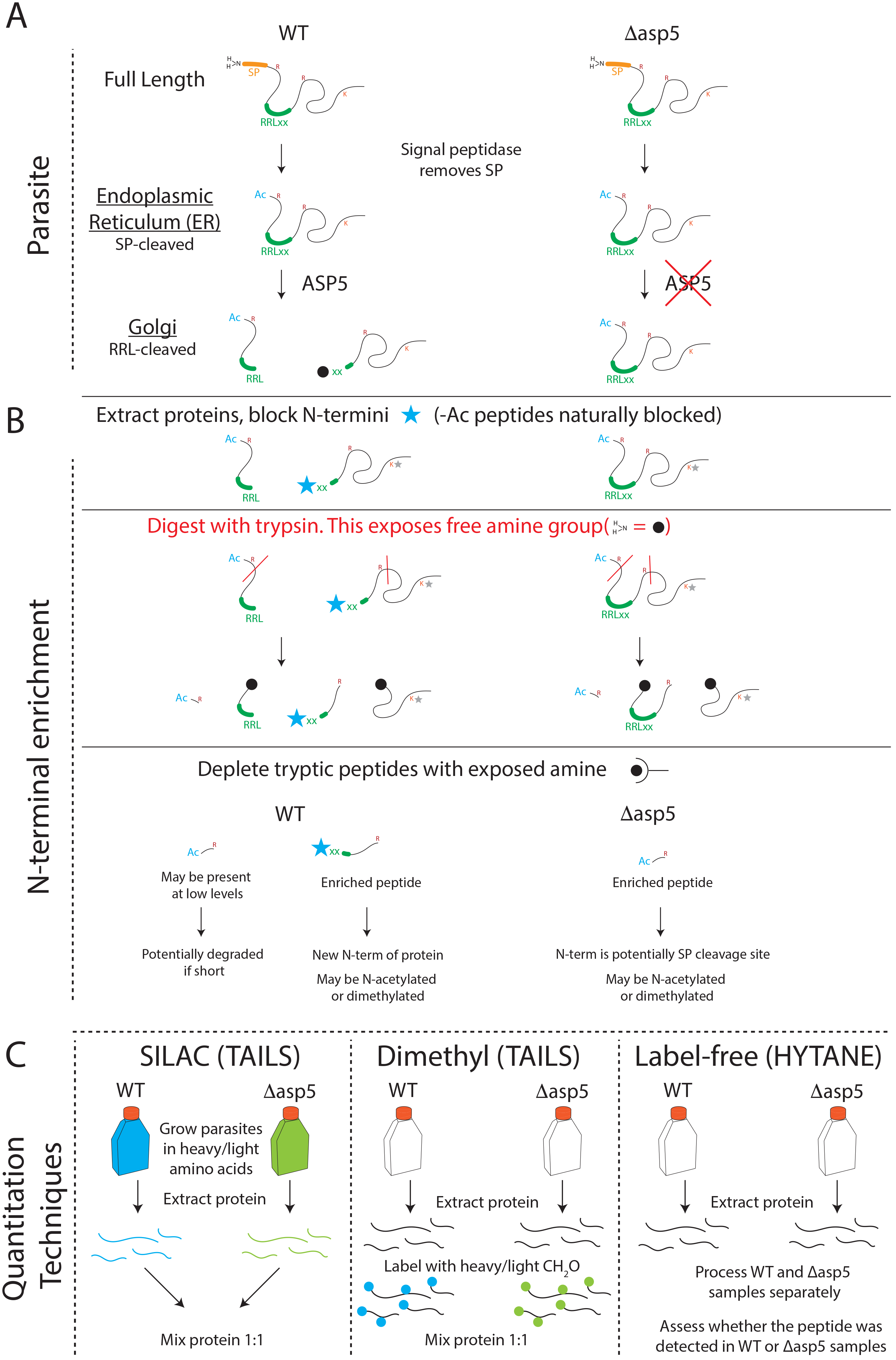
Outline of experimental techniques for identification of N-terminal peptides. (A) Schematic of TEXEL protein processing within WT (left) and Δ*asp5* (right) parasites. (B) Following protein extraction, exposed N-termini (i.e. containing a free amino group, -NH_2_, black sphere) were blocked following the addition of formaldehyde with a catalyst, resulting in primary amine dimethylation. Dimethylation of the most N-terminal amine is depicted with a blue star (-N(CH_3_)_2_), while those on the R-chain of lysine residues are depicted with a grey star. Naturally blocked N-termini, such as acetylated amines, did not undergo this reaction. The blocked N-terminal peptides were then liberated from their respective proteins by digestion with trypsin. Resulting tryptic peptides (containing an exposed NH_2_, black sphere) were then depleted from the mixture either using the synthetic HPG-ALD polymer (TAILS) or using hexadecanal (HYTANE), while the blocked peptides remained in solution; these were then desalted and run on the mass spectrometer. (C) Quantitation techniques: SILAC samples (n= 2 reciprocally labelled samples) were mixed prior to beginning the TAILS protocol, while Dimethyl samples (n= 2 reciprocally labelled samples) were mixed following heavy or light dimethylation of primary amines on day 2 of this protocol. Label-free samples were processed separately (n = 9 replicates per group), and neo-N-termini depleted with the HYTANE protocol. WT: wild type, Δ*asp5*: parasites lacking the enzyme ASP5, SP: signal peptide, NH_2_: amine group, RRLxx: TEXEL motif cleaved by ASP5, ASP5: aspartyl protease 5, Ac: acetylated N-terminus, R: arginine, K: lysine, blue star: dimethylated [N(CH_2_)_2_] N-terminal amino acid, red line: tryptic cleavage site (after R), black sphere: exposed NH_2_ group on the N-terminus, grey star: dimethylated R-chain of lysine.

To discover new ASP5 substrates we used three unbiased quantitative proteomics approaches (SILAC, heavy dimethyl labelling and label-free quantitation) in combination with two techniques to enrich N-terminally-derived peptides (TAILS and HYTANE) (Figure 1B). Both techniques rely on formaldehyde-based dimethylation to block free amines on the N-termini of proteins and side chains of lysine residues. Secondly, proteins are enzymatically digested with trypsin, liberating internal tryptic peptides (which have ‘free’ non-acetylated N-termini) from N-terminal peptides with blocked amines. The internal peptides are then depleted using an aldehyde-derived polymer (TAILS) or hexadecanal, an amine-reactive reagent (HYTANE), thus enriching for N-terminal peptides. Subsequent mass spectrometry analyses of these N-terminal peptides allow for quantitative differences to be measured in an unbiased fashion thus enabling the identification of protease cleavage sites when used in combination with protease-deficient genetic mutants.

### Quantitative proteomic approaches

We used SILAC, differential dimethylation and label-free methodologies to quantitate N-terminally-derived peptide abundance between wildtype and ASP5-deficient parasites. Prior to beginning the TAILS protocol, reciprocally-labelled SILAC proteins were isolated from WT or *Δasp5* parasites and mixed in equal concentrations (Figure 1C). The SILAC-TAILS samples were subjected to high pH fractionation to decrease sample complexity resulting in 12 total fractions per SILAC pair. For Dimethyl labelling, proteins were extracted from WT or Δ*asp5* parasites, reciprocally-blocked with either light (normal) or deuterated (heavy) formaldehyde, then mixed and processed together for the remainder of the protocol (Figure 1C). All samples were then desalted and subsequently run on the mass spectrometer to determine the identity of peptides. Relative peptide quantification with heavy dimethylation and SILAC-based strategies were performed in MS1 mode whilst area-under-the-curve measurements were performed for label-free quantitation. The N-termini in the SILAC and heavy dimethyl experiments were enriched using the TAILS method whilst the HYTANE method was used for the label-free quantitative proteomics experiments (Figure 1C).

First, we verified N-terminal enrichment using the HPG-ALD synthetic polymer previously developed for the TAILS protocol (38) (Figure 2A). In pre-TAILS SILAC samples, we found that only ~9% of all identified *Toxoplasma* peptides matched to N-terminally blocked peptides. However, upon TAILS enrichment the blocked peptides consisted of ~70% of the sample, reflecting a 7-fold N-terminal enrichment. The bulk of these modified peptides were experimentally dimethylated or naturally acetylated. Overall, we identified 2246 N-terminal peptides across the three experiments with the majority identified in the SILAC-TAILS (1505 peptides) followed by the HYTANE strategy (916 peptides) and heavy dimethylation-TAILS (327). The majority of modified peptides in the HYTANE and dimethylation-TAILS experiments were acetylated representing natural N-termini whilst the majority of peptides in the SILAC-TAILS were experimentally dimethylated (Figure 2B). TAILS should deplete tryptic peptides indiscriminately, and therefore it is not understood why there is variation in the identity of blocked peptides between these depletion methods. Using an alternative method for N-termini enrichment (HYTANE), we identified a substantial improvement in the number of modified N-termini with less than 15% being unmodified (Figure 2C). Comparison of the data sets revealed 79 peptides with modified N-termini that were common to each of the three quantitative proteomics experiments (Figure 2D). Collectively, SILAC-TAILS identified the majority of N-terminal peptides; however, each approach revealed novel peptides that were not found in the other experiments, demonstrating the importance of employing multiple N-terminome methodologies.

**Figure 2:**
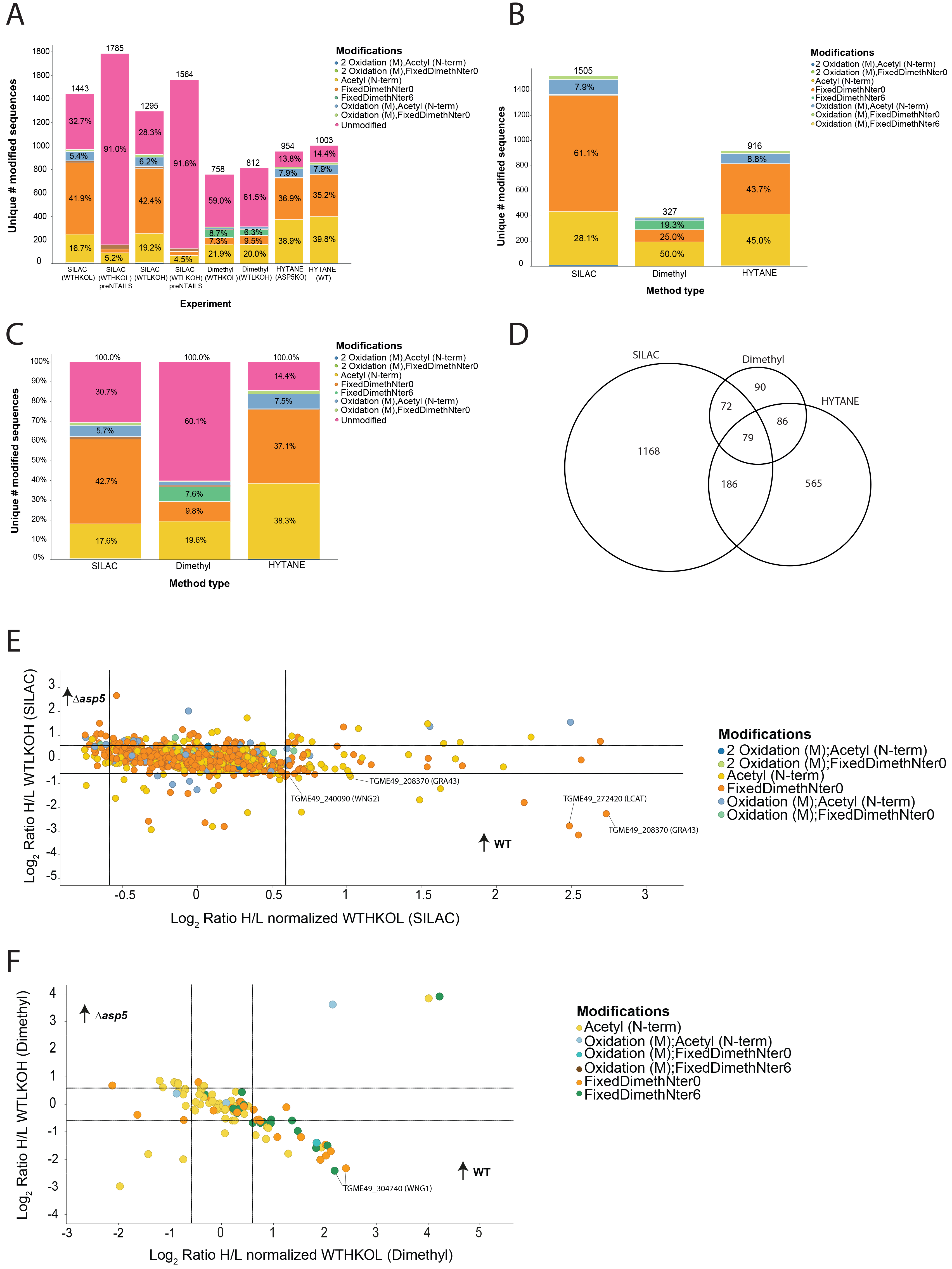
Comparison of shotgun-like preTAILS versus TAILS N-terminomics on *Toxoplasma* peptides. (A) Greater than 7-fold enrichment for N-terminal modified peptides was achieved in the SILAC samples when compared to the pre-TAILS analyses. (B) Unraveling the features of blocked N termini. The vast majority of blocked N-termini are naturally acetylated in the Dimethyl and HYTANE samples or experimentally dimethylated during TAILS in the case of the SILAC samples. (C) Overall, the HYTANE method had the lowest proportion of unmodified peptides following TAILS enrichment compared with the Overall lab method which uses the HPG-ALD polymer. (D) We identified 79 modified peptides common to all three experiments, with the majority of peptides identified uniquely to the SILAC or HYTANE experiments. (E) Log-Log plot representing the modified peptides with reciprocal protein expression from the SILAC-TAILS experiments. Peptides were deemed significantly differentially enriched in the WT sample if their Log2 ratio was above 0.59 (equivalent to 1.5 fold). (F) Log-Log plot representing the modified peptides with reciprocal protein expression from the heavy demethylation-TAILS experiments. Peptides were deemed significantly differentially enriched in the WT sample if their Log2 ratio was above 0.59 (equivalent to 1.5 fold).

We then interrogated differences in abundance of N-terminal peptides from WT and Δ*asp5* tachyzoites across all methodologies. Here, we found that the SILAC-based peptide quantitation revealed 51 *Toxoplasma* peptides with modified N-termini that were significantly differentially abundant between the WT and Δ*asp5* samples (Table S1), including three novel ASP5 substrates - TGME49_272420 (LCAT), TGME49_208370 (GRA43) and TGME49_240090 (WNG2, previously annotated as ROP34, Figure 2E). Differential dimethylation-TAILS revealed a total of 26 *Toxoplasma* peptides with modified N-termini that were significantly differentially abundant between the WT and Δ*asp5* samples (Table S1), including TGME49_304740 (WNG1, previously annotated as ROP35) (Figure 2F). Label-free quantitative proteomics analysis revealed a number of modified peptides uniquely present in the WT samples including TGME49_228170 (GRA44) and WNG2 (ROP34) (Table 1 and Table S1).

**Table 1:** List of peptides found in the combined TAILS and HYTANE datasets and subsequently validated as ASP5 substrates. Location of the peptide and three preceding amino acids were obtained from Toxodb.org v34. (fi) = fixed dimethylation from SILAC experiments; (n) = fixed dimethylation in HYTANE experiment; (Ac) = acetylation occurring within parasites and/or host cell. The HYTANE experiment did not use differential labelling so we could not directly compare ratios between WT vs Δ*asp5*; therefore, results are displayed as number of samples the peptide was detected in per condition, n = 9 (3 independent biological samples, performed in triplicate, with the HYTANE procedure performed once to reduce variation). ^a^We were unable to determine which samples (WT or Δ*asp5*) the RRL-cleaved peptide [(ac)AAEGGSESEDEQGVAR] originated from, as this was only found in the Dimethyl experiment and contains no differential heavy/light dimethylation, as the N-terminus is blocked and no there are no lysine residues. ^b^Peptide for TGME49_286530 found in both WT and Δ*asp5* parasites and maps within predicted SP, suggesting this processing is mediated by signal peptidase. ^c^We have annotated the peptide mapping to WNG1 [(Ac)TVAAPQVETGPLLSVR] as potentially SP cleaved, however SignalP 4.1 (54) does not recognise a signal peptide within this protein. This annotation is based on the peptide being N-acetylated, a modification observed predominantly at the initiator methionine, SP cleavage site and the ASP5 cleavage site. Spectra for HYTANE peptides can be found in Figure S5.

| Found following RRL (validated) |  |  |  |  |  |  |
| --- | --- | --- | --- | --- | --- | --- |
| Gene name | Gene ID<br>(TGME49_) | Sequence<br>Preceding<br>(and<br>location) | Peptide found | Up in<br>WT or<br>$\Delta asp5$? | Experiment<br>found | Ratio of<br>WT/$\Delta asp5$<br>or if<br>HYTANE,<br># samples<br>detected in |
| LCAT | 272420 | RRL<sup>476-478</sup> | (fi)DAVLTDEVGGPESGAR | WT | SILAC<br>(TAILS) | 5.61 / 0.14 |
| GRA43 | 208370 | RRL<sup>894-896</sup> | (fi)LSSSAILTGQQIGTYR | WT | SILAC<br>(TAILS) | 6.65 / 0.21 |
| GRA43 | 208370 | RRL<sup>894-896</sup> | (n)LSSSAILTGQQIGTYR | WT | LFQ<br>(HYTANE) | 8/9 WT,<br>0/9 $\Delta asp5$ |
| GRA44<br>(IMC2A) | 228170 | RRL<sup>83-85</sup> | (Ac)SGIIKTLVLWDPVQR | WT | LFQ<br>(HYTANE) | 4/9 WT,<br>0/9 $\Delta asp5$ |
| WNG2<br>(ROP34) | 240090 | RRL<sup>109-111</sup> | (n)DSLIPGFLKR | WT | LFQ<br>(HYTANE) | 6/9 WT,<br>0/9 $\Delta asp5$ |
| IST | 240060 | RRL<sup>137-139</sup> | _(ac)AAEGGSESEDEQGVAR | No<br>ratio<sup>a</sup> | Dimethyl<br>(TAILS) | Unable to<br>determine* |
| Found following RRL (not validated) |  |  |  |  |  |  |
| Hypothetical | 233695 | RRL<sup>115-117</sup> | QAGVYFSEEDR | WT | LFQ<br>(HYTANE) | 2/9 WT,<br>0/9 $\Delta asp5$ |
| Hypothetical | 297890 | RRL<sup>183-185</sup> | (n)TTLASTLSLSR | WT | LFQ<br>(HYTANE) | 2/9 WT,<br>0/9 $\Delta asp5$ |
| Hypothetical<br>(Zinc<br>Finger-<br>containing) | 248450 | RRL<sup>345-347</sup> | (Ac)YAPGASVVESPVFGTPPSR | WT | LFQ<br>(HYTANE) | 1/9 WT,<br>0/9 $\Delta asp5$ |
| Hypothetical | 286530 | RRL<sup>27-29</sup><br>SP-cleaved<sup>b</sup> | (n)MFAAAPLQSFSVTNKQFHPE<br>GLEAQAPRPHQGLDMR | Neither | LFQ<br>(HYTANE) | 4/9 WT,<br>2/9 $\Delta asp5$ |

| Found following predicted signal peptide cleavage site |  |  |  |  |  |  |
| --- | --- | --- | --- | --- | --- | --- |
| WNG2<br>(ROP34) | 240090 | AVA<sup>62-64</sup><br>(SP<br>cleaved) | (n)AHAHPEDSATNFLSFAENS<br>LANREPPEDSAARPSSR | $\Delta asp5$ | LFQ<br>(HYTANE) | 0/9 WT,<br>7/9 $\Delta asp5$ |
| WNG2<br>(ROP34) | 240090 | AVA<sup>61-64</sup><br>(SP<br>cleaved) | (fi)AHAHPEDSATNFLSFAE<br>NSLANR | Neither | SILAC<br>(TAILS) | WTH/KOL<br>1.49 / 0.67<br>(not<br>significant) |
| WNG1<br>(ROP35) | 304740 | AGA<sup>68-70</sup><br>(SP<br>cleaved?)<sup>c</sup> | (Ac)TVAAPQVETGPLLSVR | WT | Dimethyl<br>(TAILS) | WT Light<br>DiMethyl<br>5.31 / 0.18 |
|  |  |  |  |  |  | WT Heavy<br>DiMethyl<br>4.17 / 0.81 |

In total, we identified over 2000 unique N-terminal peptides across the three methodologies. Many of these peptides were not significantly different between WT and Δ*asp5* parasites, having arisen from natural N-termini (both exposed and acetylated) as well as from other protease cleavage events within parasites. However, each methodology also revealed unique peptides that mapped directly after an RRL motif, enabling the identification of likely novel ASP5 substrates.

### The PVM protein LCAT is processed by ASP5

We sought to validate newly identified proteins as ASP5 substrates. To do this we endogenously tagged candidate proteins in both wildtype and *Δasp5* tachyzoites, then mutated the putative RRL cleavage site to look for differential processing within parasites. For tagging and mutagenesis of almost all proteins, we designed Cas9-targeting guides against the gene-of-interest and co-transfected these with two annealed ~60 base pair oligos to introduce either an epitope tag, mutate an RRL⟶ARL, or disrupt the gene (Figure 3A). We first chose to investigate a recently discovered dense granule protein LCAT (TGME49_272420) that is secreted to the PVM, as we identified an N-terminal peptide found exclusively in WT parasites with the sequence (dimethyl)-DAVLTDEVGGPESGAR (Table 1), which mapped to a location directly C-terminal of an RRL sequence within this protein (40). LCAT is processed into two fragments by an unknown protease within the ‘inserted element (IE)’ that separates the catalytic residues of the enzyme (40). As the TAILS peptide mapped to directly after an RRL within this IE, we tagged endogenous LCAT and observed a ~90 kDa ‘full-length’ species and a ~40 kDa processed form (Figure 3B). We then tagged LCAT in Δ*asp5* parasites and primarily observed the larger molecular weight species (Figure 3B), with a doublet band at ~60 kDa that may be a degradation product or result from a subsequent processing event. To confirm that the loss of the ~40 kDa species was mediated by ASP5, we then swapped the endogenous RRL to ARL (schematic in Figure 3A) and observed only the larger species in WT LCAT_ARL_-HA. Together, this strongly suggests that LCAT is processed at the TEXEL motif identified by TAILS (Table 1).

**Figure 3;.**
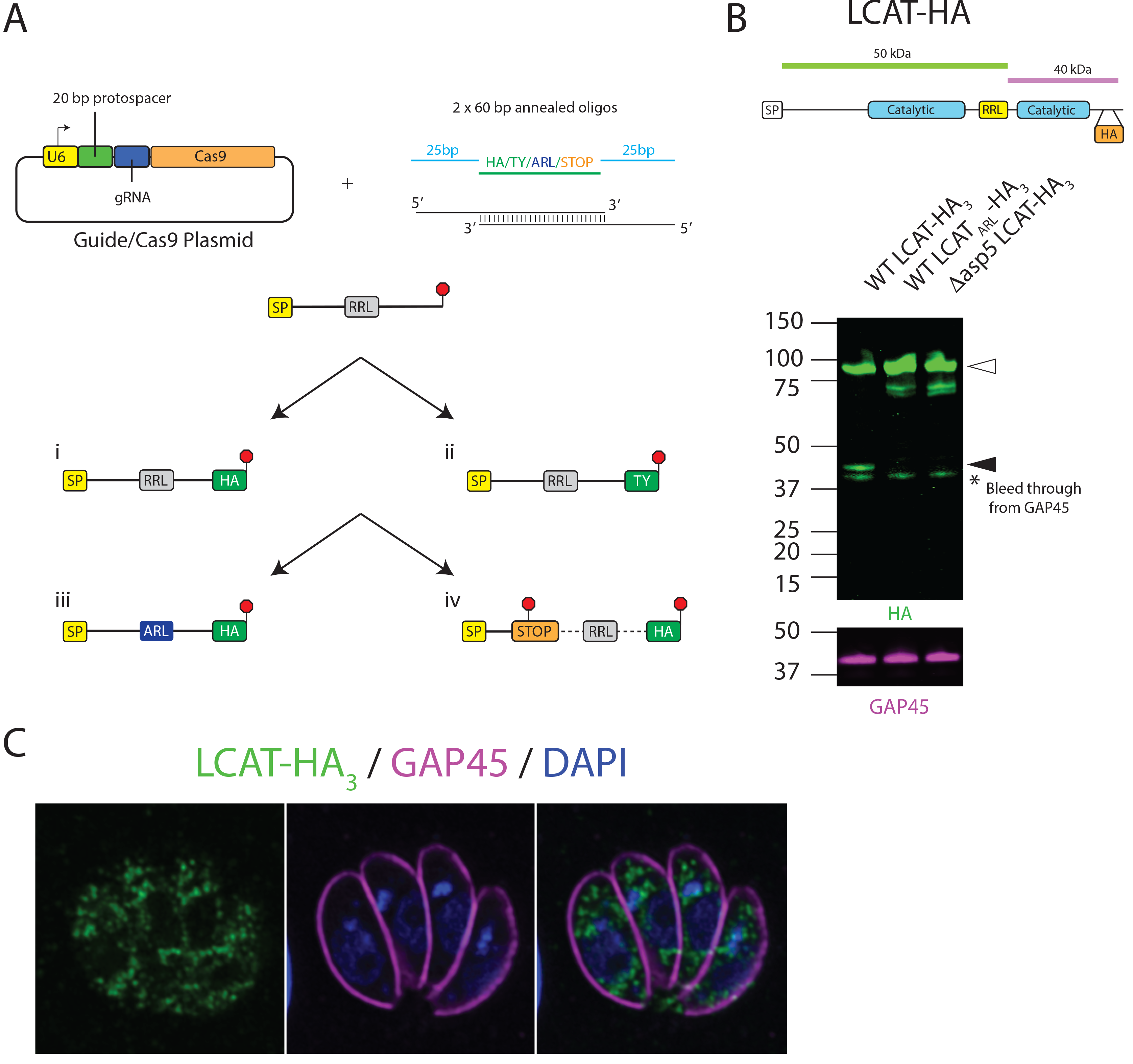
Methodology for validation of ASP5 substrates. (A) Schematic of endogenous tagging, U6: U6 promoter, protospacer: 20-21 base pair sequence used to direct Cas9 to cut within the parasite genome, gRNA: guide RNA, Cas9: enzyme that cuts DNA, SP: signal peptide, RRL: arginine; arginine; leucine, site of cleavage by ASP5, HA: haemagglutinin tag, Ty: Ty tag, Red stop signal: endogenous stop codon or introduced stop codon/frameshift mutation, ARL: mutation of first arginine to alanine. (B) Immunoblot of LCAT-HA3 (TGGT1_272420) using the LI-COR Odyssey imager, αHA and aGAP45 antibodies used. Open arrow heads indicate full length species, closed arrow heads indicate ASP5-cleaved species. (C) IFA of LCAT-HA_3_ expressing parasites, HA and GAP45 antibodies used (LCAT: TGGT1_272420).

It is important to note that by immunofluorescence assays (IFA), we could only detect LCAT within punctate structures within parasites, possibly the dense granules (Fig 3C), but not at the PVM as has previously been reported (40). The absence of signal at the PVM suggests that the introduction of the 3xHA tags or replacement of the endogenous 3’ UTR with the DHFR 3’ UTR interfered with trafficking of this enzyme (Figure 3C). To overcome this, we attempted to introduce a single HA tag into the endogenous locus as this has been shown not to affect trafficking (40). However, despite several attempts we were unable to introduce this single HA-tag into the endogenous LCAT locus without positive selection, and therefore we were unable to assess any differences in localisation in LCAT between WT and Δ*asp5* parasites (data not shown).

### N-terminal peptide enrichment identifies novel dense granule proteins as ASP5 substrates

After validating the dense granule protein LCAT as an ASP5 substrate, we sought to investigate novel and hypothetical candidates. One peptide arising after an RRL only in WT parasites was Ac-SSSAILTGQQIGTYR (Table 1), mapping to a hypothetical protein (TGME49_208370), which we have named GRA43. This protein was chosen for further investigation as the RRL motif maps near the C-terminus of the protein prior to a predicted transmembrane domain, similar to MYR1 (Figure 4A). GRA43 is annotated on ToxoDB as ‘myosin heavy chain-like’, but further investigation has revealed limited homology to myosin, rendering it unlikely that this protein is a true myosin (data not shown). To assess GRA43 as a potential ASP5 substrate, we inserted a HA tag at the C-terminus in WT parasites and observed two species, one at approximately 22 kDa and another at ~ 30 kDa, both well below the predicted size of ~125 kDa (Figure 4A and B). We then repeated this process in Δ*asp5* parasites, revealing the lowermost band and a second at ~140 kDa, indicating the loss of a processing event, dependent on this aspartyl protease. To validate the RRL found by TAILS within this vicinity, we mutated the RRL⟶ARL (Figure S1A) and again observed the same processing pattern as seen in Δ*asp5* parasites, strongly suggesting that GRA43 is cleaved by ASP5 in a TEXEL-dependent manner. The presence of the lowest molecular weight species in WT, ARL and Δ*asp5* parasites potentially results from processing at or near the predicted transmembrane (TM) domain near the C-terminus of this protein by a protease other than ASP5 (Figure 4A and B).

**Figure 4:**
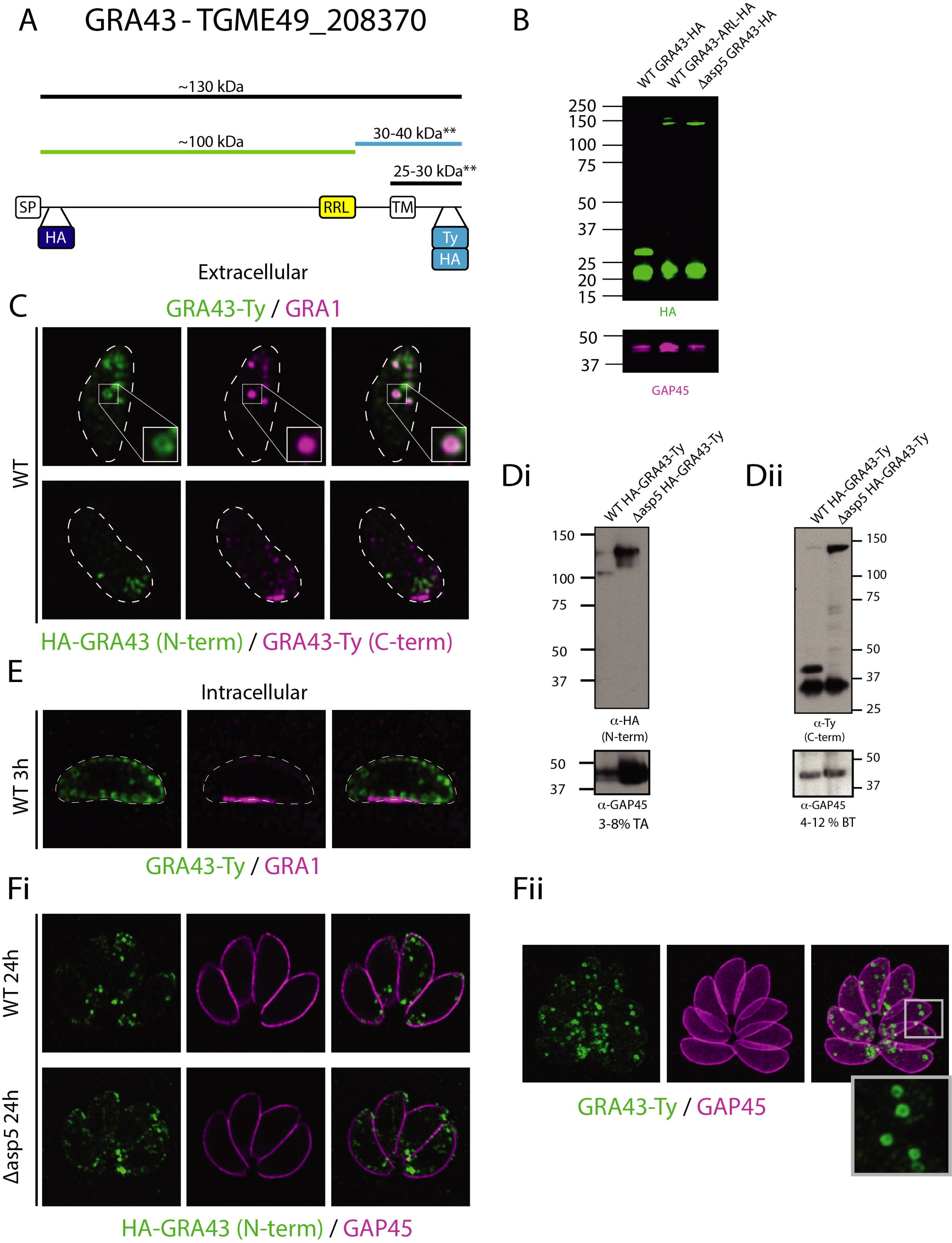
GRA43 is a novel ASP5 substrate. (A) Schematic of TGGT1_208370 (GRA43) with a HA tag inserted after the signal peptide cleavage site, a TEXEL (RRL), transmembrane domain (TM) and TY or HA tag inserted prior to the stop codon. (B) i – immunoblot of 3-8 % Tris-Acetate (TA) gel using αHA antibodies (N-terminal fragment), ii – using αTy antibodies (C-terminal fragment). Open arrow heads indicate full length species, closed arrow heads indicate ASP5-cleaved species. (C) Extracellular IFA. (D) Immunoblot using the LI-COR Odyssey imager. (E) IFA at 3 hours post invasion. (F) IFA at 24 hours post invasion of replicating parasites.

As both MYR1 and LCAT are cleaved by ASP5 near the C-terminus, and then the N- and C-termini re-associate (37, 40), we sought to determine whether the same was true for GRA43. To address this, we tagged GRA43 at the C-terminus with a TY tag, followed by a HA tag shortly after the predicted signal peptidase cleavage site (Figure 4A). To determine the sub-cellular localisation of the two ASP5-processed sequences of GRA43, we probed extracellular parasites with αTy antibodies using super-resolution microscopy and observe a spherical donut-shaped localisation that encloses GRA1 staining in dense granules, suggesting this protein resides at the periphery of the dense granules (Figure 4C). We observe very low levels of co-localisation of the N-terminal (αHA) and C-terminal (αTy) fragments of GRA43 in extracellular parasites (Figure 4C, bottom panel).

Immunoblot analysis of WT parasites expressing HA-GRA43-TY using αHA antibodies revealed two bands at approximately 130 and 100 kDa, while only the higher molecular weight species was present in *Δasp5* parasites, suggesting a loss of processing (Fig 4Di). Using αTy antibodies the majority of this protein was detected at ~40 kDa, while in *Δasp5* parasites this species was lost and the protein remained at a higher molecular weight, consistent with the full-length protein (Figure 4Dii). As in Figure 4B, in both WT and *Δasp5* parasites we observed a consistent ~ 30 kDa band for GRA43 that is consistent with processing around the TM domain of this protein by a protease other than ASP5 (Figure 4Dii).

We then proceeded with further assessment of the localisation of GRA43. To determine the location of this protein during early infection, parasites were fixed at 3 hours post invasion, which revealed the secretion of GRA1 into the PV, but GRA43 appeared to remain associated with the dense granules (Figure 4E). At 24 hours post invasion, the N-terminal processed species of GRA43 in WT and Δ*asp5* parasites localised to discrete puncta within the parasites, notably at the anterior and posterior ends of the tachyzoites (Figure 4Fi). At this time point, the C-terminal fragment also localised to similar spherical structures within the parasites, potentially dense granules (Figure 4Fii). We could detect no difference in localisation of the C-terminal fragment between WT and Δ*asp5* parasites.

Together, these data reveal that GRA43 is processed by ASP5 near the C-terminus, and that it is a dense granule protein based on the localisation of both cleaved fragments. The TEXEL of GRA43 maps to the C-terminal end of this protein; however, unlike MYR1 and LCAT that are cleaved by ASP5 within a similar region, the N- and C-terminal fragments of GRA43 do not appear to re-associate. Similar to MYR1 and LCAT, we were unable to detect GRA43 within the host cell, suggesting that it is not exported and raising the possibility that processing near the C-terminal region could be a marker of non-exported ASP5 substrates.

### Two WNG kinases are substrates of ASP5

Further investigation of our proteomic dataset revealed two proteins that were previously annotated as rhoptry kinases through their similarity to ROP16, ROP18 and several other predicted kinases (Table S1) (41). Here we present ROP35 (TGME49_304740) and ROP34 (TGME49_240090), herein named WNG1 and WNG2 respectively for reasons outlined below, as novel ASP5 substrates. WNG1 was identified in the TAILS screen due to increased levels of acetylated SP-cleaved peptide within Δ*asp5* parasites (Table 1), that mapped upstream of a TEXEL motif, suggesting that the SP-cleavage site is the N-terminus in Δ*asp5* parasites.

To determine the fate of WNG1, we epitope-tagged this protein at the C-terminus and monitored its maturation by western blot (Figure 5A and B). We observed that WNG1 expressed by WT parasites was present at ~42 kDa, which is a smaller size than predicted by the full amino acid sequence (Figure 5A). In contrast, epitope tagging in Δ*asp5* parasites revealed that WNG1 was present at a higher molecular weight, suggesting it is indeed processed by ASP5 (Figure 5B). To assess whether WNG1 is processed within the predicted TEXEL motif, we mutated the endogenous RRL ⟶ ARL (Figure S1B), and indeed, observed an accumulation of the higher molecular weight species in these WNG1_ARL_-HA parasites (Figure 5B).

**Figure 5:**
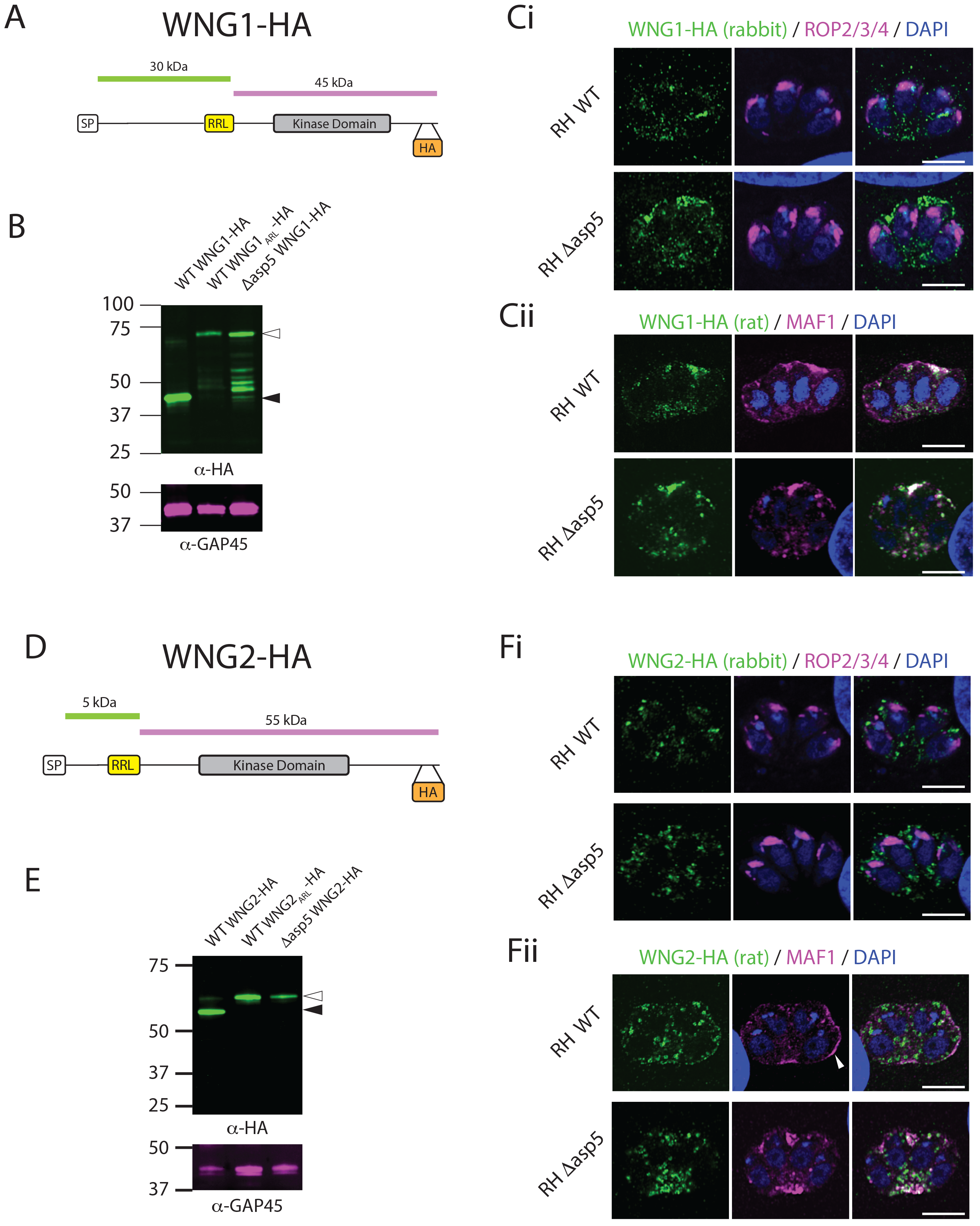
WNG1 and WNG2 are ASP5 substrates that localise to the host-parasite boundary. (A) Schematic of WNG1. (B) Immunoblot using the LI-COR Odyssey imager. Open arrow heads indicate full length species, closed arrow heads indicate ASP5-cleaved species. (C) IFAs at 24 hours post invasion with rhoptry markers αROP2/3/4 or the dense granule marker MAF1. (D) Schematic of WNG2. (E) Immunoblot using the LI-COR Odyssey imager. Open arrow heads indicate full length species, closed arrow heads indicate ASP5-cleaved species. (F) IFAs at 24 hours post invasion with rhoptry markers αROP2/3/4 or the dense granule marker MAF1.

We then investigated the localisation of WNG1 by IFA and found no co-localisation with the αROP2/3/4 marker in intracellular parasites, suggesting this predicted kinase is not a true rhoptry protein (Figure 5Ci). The low level of HA signal observed using rabbit αHA antibodies in WT WNG1-HA expressing parasites (Figure 5Ci, upper panel) was more pronounced in Δ*asp5* parasites (lower panel), suggesting that loss of ASP5 leads to the accumulation of WNG1 within localised areas in the PV, a phenomenon observed for several dense granule proteins following deletion of ASP5 (20, 22) (Figure 5Ci). WT:WNG1-HA was primarily found within the vacuole and exhibited partial co-localisation with the PV/PVM marker MAF1 (Figure 5Cii), as did Δ*asp5*:WNG1-HA, suggesting WNG1 is a dense granule protein.

We then investigated WNG2 and its dependence on ASP5 for proteolytic maturation. Our N-terminome data identified a peptide that mapped to WNG2 resulting from cleavage after an RRL (dimethyl-DSLIPGFLKR) was observed in WT but not in Δ*asp5* parasites (Table 1). To confirm if WNG2 is a true ASP5 substrate, we inserted a HA tag into the endogenous locus just before the stop codon and observed the dominant species at ~55 kDa, despite the predicted molecular weight of ~62 kDa. In contrast, in parasites lacking ASP5, and those expressing the mutated RRL⟶ARL (WNG2_ARL_-HA, Figure S1D), this protein migrated at approximately 60 kDa, strongly suggesting ASP5-dependent maturation (Figure 5E).

We then investigated the localisation of WNG2 by IFA. As with WNG1, WNG2 did not co-localise with the rhoptry marker αROP2/3/4 (Figure 5Fi). When probing with antibodies against the PV/PVM protein MAF1 (Figure 5Fii), a small proportion of WNG2-HA expressed by WT parasites co-localised at the PVM, however most was present within the parasites within the donut-shaped spheres, similar to that described for GRA43, suggestive of dense granules (Figure 4C). In Δ*asp5* parasites, MAF1 loses a significant proportion of its PVM localisation, as has been previously reported (20), and the WNG2-HA signal in Δ*asp5* parasites overlaps with the MAF1 signal detected in the PV space. Our data therefore strongly suggests that these kinases are unlikely to represent bona fide rhoptry proteins and we have therefore renamed these proteins as WNG kinases, short for ‘With No Gly-loop’ as they do not contain the characteristic glycine-rich loop of other kinases. As ROP35 is the most conserved across Coccidia, we named this WNG1, with ROP34 renamed as WNG2.

### ASP5 matures the dense granule phosphatase GRA44 and a novel dense granule protein

Our N-terminome analyses discovered a peptide resulting from cleavage after an RRL within the protein annotated as inner membrane complex protein 2A (IMC2A, TGME49_228170). For reasons outlined below, we rename IMC2A to GRA44.

To investigate the maturation of GRA44 by ASP5 we introduced a HA tag shortly before the endogenous stop codon in WT parasites (Figure 6A). We observed a species of ~37 kDa, shorter than the predicted ~185 kDa size of the full-length protein (Figure 6A and B). In contrast, this protein migrated close to 200 kDa in Δ*asp5* parasites, suggesting the loss of one or more processing events within these parasites. The RRL-cleaved peptide of GRA44, found in both the dimethyl and HYTANE experiments, was observed only in WT parasites and maps near the predicted SP-cleavage site. However, we were unable to mutate this endogenous RRL⟶ARL despite several attempts, suggesting a fitness cost to the parasites (data not shown). Processing at this N-terminal RRL could not account for the ~37 kDa species observed in WT parasites, suggesting GRA44 is processed at least twice. Indeed, GRA44 contains a second RRL approximately 32 kDa from the C-terminus (Figure 6A) which may be processed by ASP5.

**Figure 6:**
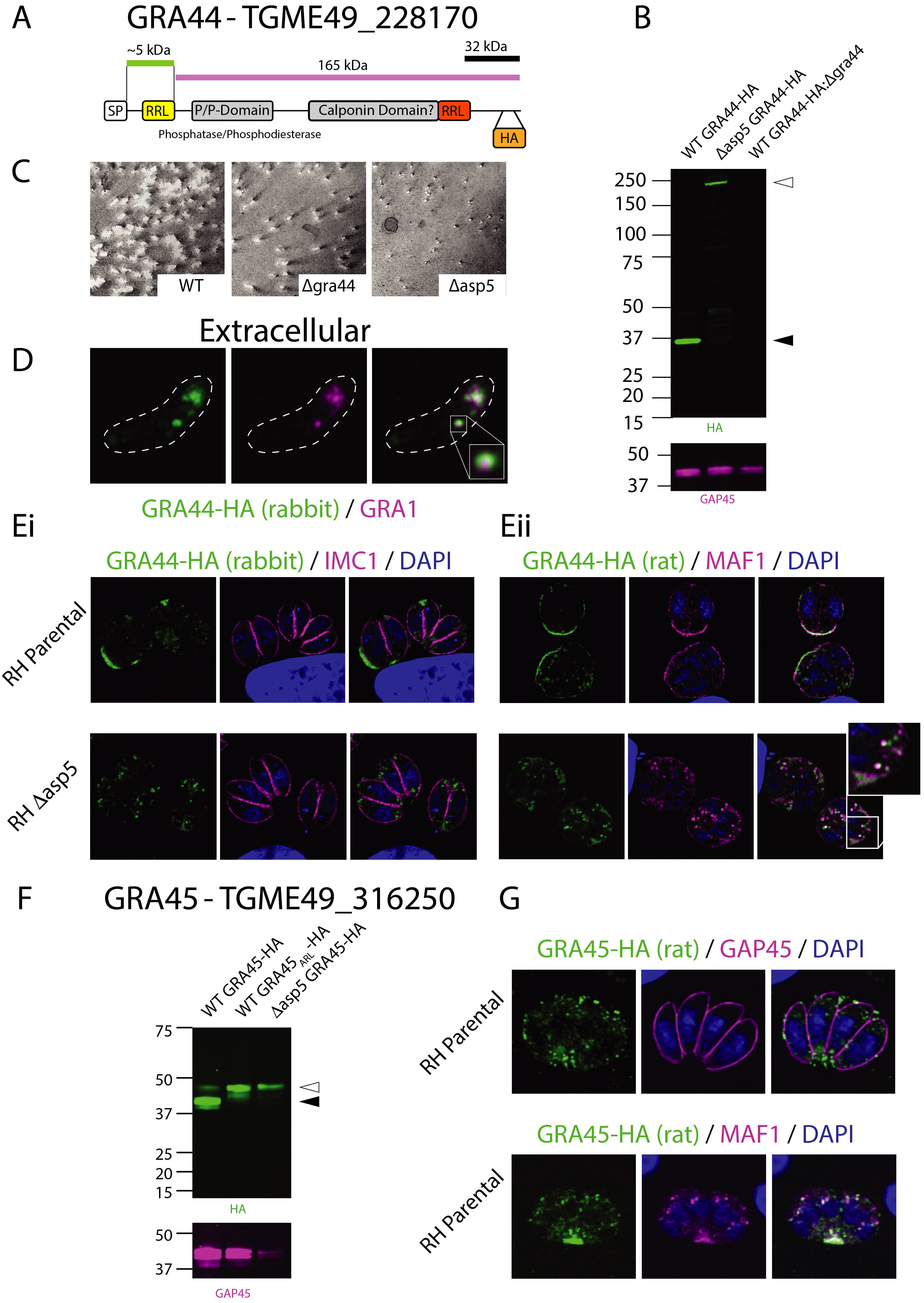
GRA44 is matured by ASP5 and localises to the PVM. (A) Schematic of GRA44 based off the TGGT1_228170 sequence. (B) Immunoblot using the LI-COR Odyssey imager, αHA and αGAP45 antibodies used. Open arrow heads indicate expected size of full length species, closed arrow heads indicate predicted ASP5-cleaved species. (C) Plaque assay at 9 days depicting plaque sizes for WT (RHΔ*ku80*Δ*hx*), Δ*gra44* (RHΔ*ku80*Δ*hx*Δ*gra44*) and Δ*asp5* (RHΔ*ku80*Δ*hx*Δ*asp5*). (D) IFA of an extracellular tachyzoite depicting WT GRA44-HA (green) and the dense granule marker GRA1 (magenta). (E) Intracellular IFAs of WT- and Δ*asp5*-GRA44-HA parasites with the IMC marker IMC1 (E-i) and the dense granule marker MAF1 (E-ii). (F) Immunoblot using the LI-COR Odyssey imager, αHA and αGAP45 antibodies used. Open arrow heads indicate the predicted full-length species, closed arrow heads indicate predicted ASP5-cleaved species. (G) IFA of WT intracellular parasites expressing GRA45-HA (green) with GAP45 (upper panel) and MAF1 (lower panel).

To investigate the importance of GRA44 on parasite growth we generated a knockout line by integration of CAT selectable marker (Figure S1D), which led to the ablation of HA signal by western blot (Figure 6B, longer exposure in Figure S3). Parasites lacking GRA44 (Δ*gra44*) formed smaller plaques than WT parasites, yet the defect was not as severe as that observed for Δ*asp5* parasites, despite having a lower CRISPR fitness score (42) (Figure 6C). In extracellular parasites, GRA44 -HA signal enclosed a subset of GRA1-containing dense granules, similar to GRA43 (Figure 6D, 4C). Immunofluorescence assays demonstrate a lack of co-localisation with IMC1 in intracellular parasites, suggesting this protein is not part of the IMC, as its previous annotation of IMC2A suggests, and instead localises within the PV and PVM in WT parasites (Figure 6Ei, ii). In *Δasp5* parasites this protein is observed primarily between parasites in the PV instead of the PVM, similar to MAF1 (Figure 6E), as is common for dense granule proteins after deletion of ASP5 (20–22).

To further elucidate the mechanisms by which the novel ASP5 substrates operate, we performed immunoprecipitations to assess whether any act within larger complexes. To address this, we infected HFFs at a MOI of 5 with GRA43-HA, GRA44-HA and WNG2-HA parasites, then harvested from large vacuoles at 36 hours post infection, lysed and then pulled down these proteins using aHA antibodies. IPs were performed in triplicate and protein eluates were subjected to mass spectrometry analysis where protein expression changes were quantified using a custom label-free pipeline. Each of the pulldowns enriched the bait (Figure S2), while also enriching for several other dense granule proteins. TGME49_316250 (GRA45) was highly enriched in GRA44-HA and WNG2-HA pulldowns compared to GRA43-HA (Supp. Fig 2) and was chosen for further validation as it contains a signal peptide and a putative TEXEL motif.

To investigate TGME49_316250, which we herein rename GRA45, we epitope tagged this protein just before the predicted stop codon. The dominant molecular weight species observed in WT parasites was approximately 42 kDa, with a fainter band at ~47 kDa (Figure 6F). Deletion of ASP5, or mutation of the RRL⟶ARL resulted in the loss of the lower migrating band, suggesting this protein is indeed processed by ASP5 (Figure 6F). GRA45-HA was observed in punctate structures within parasites and the PV space, primarily at the posterior of parasites (Fig 6G), somewhat overlapping with the dense granule protein MAF1. Similar localisation was observed in Δ*asp5* parasites (not shown), suggesting this novel ASP5 substrate is a dense granule protein.

### WNG2 and GRA43 are important for virulence

ROP kinases are important virulence factors (3–11), therefore we wanted to determine if the WNG kinases also played important roles *in vivo*. We generated knockouts of WNG1, WNG2 and GRA43 (Figure S1A, B and C) and confirmed loss of expression using western blot (Figure 7A, B and C). Furthermore, we generated knockout of ASP5 in a type II Pru background for purposes of comparison (Figure S1E). As has been previously reported, murine infection with less than 100 RH*Δasp5* parasites leads to attenuation in virulence (20, 22), however higher doses render mice susceptible in a similar manner to infection with WT parasites. To confirm if this was true for Pru*Δku80* parasites, we injected six C57/BL6 mice with 2000 Pru*Δku80* (WT) or Pru*Δku80Δasp5*(*Δasp5*) parasites and observed complete attenuation in mice infected with *Δasp5*, while all mice infected with WT parasites succumbed by day 8 (Figure 7D). The same was true when infection was increased to 5,000 or 50,000 parasites (Fig 7E), with mice infected with *Δasp5* parasites retaining their initial bodyweight (lower panel), despite exhibiting bloating of the abdomen, a common response to infection.

**Figure 7:**
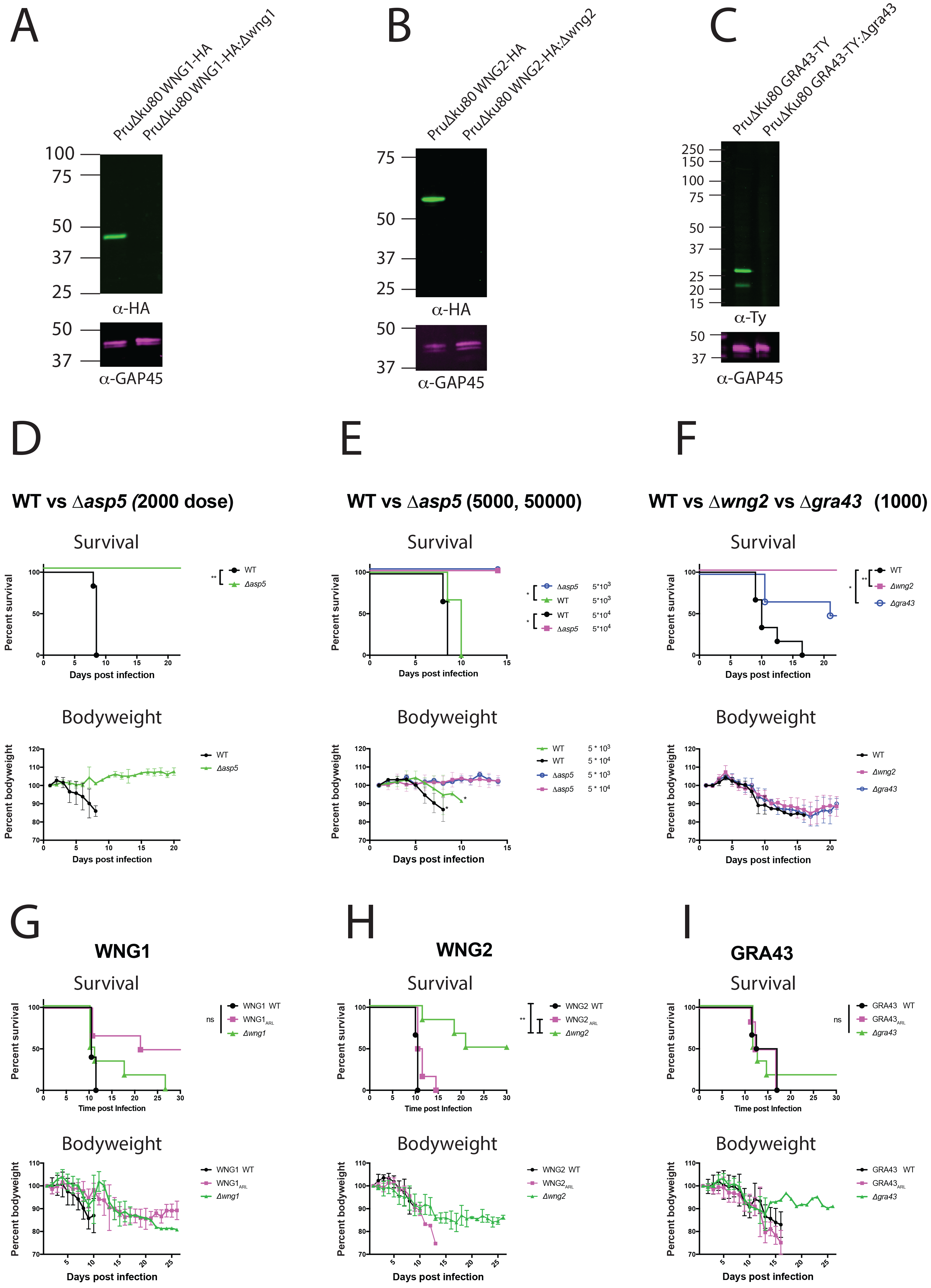
WNG2 and GRA43 are important for acute virulence. (A-C) Immunoblots confirming genetic disruption of WNG1 (A), WNG2 (B) and GRA43 (C) in PruΔ*ku80* parasites. (D-I) Kaplan-Meier survival curves of C57BL/6 mice with strains as indicated (upper panels) and pooled bodyweight (lower panels). Survival curve comparisons were assessed using the Log-Rank (Mantel-Cox) test and significance assessed following correction for the Bonferroni threshold when performing multiple comparisons. (D) Six C57BL/6 mice were infected with 2000 WT (PruΔ*ku80*) or Δasp5 (PruΔ*ku80*Δ*asp5*) tachyzoites via intraperitoneal (IP) injection. p = 0.0012. (E) C57BL/6 mice (n = 3) were infected via IP injection with 5*10^3^ or 5*10^4^ WT (PruΔ*ku80*) or Δ*asp5*(PruΔ*ku80*Δ*asp5*) tachyzoites. p = 0.0295 for both comparisons. (F) Mice (n = 6) were infected via IP injection with 10^3^ tachyzoites of WT (PruΔ*ku80*), Δ*wng2*(PruΔ*ku80*Δ*wng2*) and Δ*gra43* (PruΔ*ku80*Δ*gra43*) backgrounds. Significance: WT vs Δ*wng2*, p = 0.0012. WT vs Δ*gra43*, p = 0.0023. (G) Infection via IP injection with PruΔ*ku80* WNG1-HA (WNG1 WT), PruΔ*ku80* WNG1_ARL_-HA (WNG1 ARL) and PruΔ*ku80*Δ*wng1* (Δ*wng1*). n = 5, 6 and 6 respectively. (H) Infection via IP injection (n = 6) with PruΔ*ku80* WNG2-HA (WNG2 WT), PruΔ*ku80* WNG2_ARL_-HA (WNG2_ARL_) and PruΔ*ku80*Δ*wng2* (Δ*wng2*). Statistics: WNG2 WT vs Δ*wng2*, p = 0.0012. WNG2_ARL_ vs Δ*wng2*, p = 0.0029. (I) Infection via IP injection (n = 6) with PruΔ*ku80* GRA43-HA (GRA43 WT), PruΔ*ku80* GRA43_ARL_-HA (GRA43_ARL_) and PruΔ*ku80*Δ*gra43* (Δ*gra43*).

We then sought to determine whether GRA43 and WNG2 are important for parasite virulence during acute infection. To address this, we injected six C57/BL6 mice with 10^3^ Pru*Δku80* WT, Pru*Δku80Δgra43*(*Δgra43*) and Pru*Δku80Δwng2* (*Δwng2*) tachyzoites then monitored infection (Figure 7F). All mice infected with WT parasites gradually lost bodyweight throughout infection and were culled between days 8-17. Mice infected with *Δgra43* and *Δwng2* parasites lost bodyweight, however by day 17 the remaining mice began to recover. All mice infected with *Δwng2* parasites survived the infection, while 3/6 infected with *Δgra43* were culled. All surviving mice were seropositive for exposure to *Toxoplasma* (data not shown). Despite these defects we could not observe any growth defects *in vitro*, as monitored by plaque assay (Figure S4).

After establishing that *Δgra43* and *Δwng2* tachyzoites have a mild virulence defect *in vivo*, we sought to interrogate whether ASP5 cleavage is required for this virulence shift. To do this we used tachyzoite lines harbouring RRL⟶ARL mutations (Figure S1) that are not proteolytically processed (Figure 4B, 5B and 5E). We then infected C57/BL6 mice at a slightly higher inoculum to further dissect out virulence defects (Figure 7G-I), and simultaneously performed plaque assays, revealing all mice received the same infectious dose and there were no observable growth defects *in vitro* for these strains (Figure S4). All mice infected with the parental WNG1-HA strain were culled by day 11 post infection, while those infected with WNG1_ARL_-HA succumbed between days 10-26 (Figure 7G). Three mice infected with Δ*wng1* parasites were culled throughout the experiment, while a further three began to recover their initial bodyweight loss (Figure 7G, lower panel) and survived until day 30 post-infection, when the experiment was concluded. Differences in survival were assessed using the Log-rank (Mantel Cox) test, adjusted according to the Bonferroni correction and were not statistically significant between any of the strains.

Mice infected with the parental WNG2-HA parasite line were culled by day 10 post infection, while all mice infected with WNG2_ARL_-HA parasites were moribund by day 15 (Fig 7H). In contrast, 3/6 mice infected with Δ*wng2* parasites survived the experiment, despite stagnating at ~90 % of initial bodyweight between days 15-30 post infection.

While infection with 1000 Δ*gra43* parasites led to an attenuation of virulence, this was not replicated after the dose was doubled to 2000 parasites, suggesting this protein has a very modest effect during acute infection (Figure 7F, I). In contrast, Δ*wng2* parasites were significantly less virulent than WT parasites at both doses, indicating an important function during acute infection. Furthermore, for all TEXEL proteins assessed in this experiment, there was no statistical difference between WT and RRL⟶ARL mutants, suggesting that for these non-exported ASP5 substrates, efficient processing by ASP5 is not essential for their function during murine infection.

## Discussion

*Toxoplasma* uses a repertoire of secreted and exported proteins to exquisitely modulate the host response to infection. Initial host cell subversion is achieved through the secretion of the rhoptry proteins, notably ROP16 and ROP5/18 (3–5). Following establishment of the PV, *Toxoplasma* translocates a second wave of proteins across the PVM. These include two ASP5-cleaved exported proteins, GRA16 and IST, that *Toxoplasma* employs to regulate the cell cycle and avoid clearance from cells following IFNγ signalling, respectively. Furthermore, GRA24 does not appear to be processed by ASP5, yet its translocation across the PVM is still dependent on this protease (16, 20–22). Interestingly, the translocation of these proteins into the host cell requires the PVM protein MYR1, another ASP5 substrate. Despite the importance of this protease, only a handful of ASP5 substrates have been described to date. To address this shortcoming and identify new *Toxoplasma* effectors and virulence factors, we have exploited the differences in N-termini between WT and Δ*asp5* parasites using a quantitative proteomics approach in combination with N-terminal enrichment methods. In doing so, we have discovered several new effectors. We have validated several of these substrates, including confirmation of their processing by ASP5.

Here we have presented that WNG1, WNG2, GRA44, GRA43 and GRA45 are processed by ASP5. Notably, immunofluorescence of each of these proteins has revealed they are at least transient components of the dense granules. It is currently unclear whether ASP5 processes substrates from other organelles, including the micronemes and the rhoptries, however, proteins from these organelles have recently been demonstrated to be processed by aspartyl protease 3 (ASP3) (43), suggesting different proteases may function in maturation of proteins destined for different cellular compartments. Following ASP5 cleavage, we observe secretion of these newly validated substrates into the PV/PVM, with no signal detected within the host cell. However, we cannot preclude that some of these proteins are indeed translocated across the PVM but subsequently are highly diluted and unable to be detected through normal immunofluorescence imaging. Indeed, the four characterised exported proteins GRA16, GRA24, GRA28 and IST all traffic to the host nucleus (14, 16–19), potentially concentrating their signal above levels that could be detected if they were throughout the cytosol.

Unlike in *Plasmodium*, the TEXEL sequence in *Toxoplasma* appears not to be spatially restricted. The identification of new ASP5 substrates within this manuscript confirms that the TEXEL can be found anywhere throughout the protein. However, this motif in the two exported substrates, GRA16 and IST, is located approximately 40 and 70 amino acids from the predicted SP cleavage site, or at 13 and 16 % of the protein length respectively. As GRA44 is also processed within this approximate region, cleavage by ASP5 alone cannot dictate translocation across the PVM. It is possible ASP5 processing potentially liberates an export signal, such as the linear or structural sequence of amino acids revealed following cleavage, a suggestion that has also been raised for export of PEXEL proteins (28). To address this, both a non-exported and an exported protein should be monitored for their ability to translocate the PVM following targeted mutagenesis of the TEXEL and surrounding residues. This could reveal whether residues either upstream or downstream of cleavage are important for subsequent trafficking and ultimately export into the host cell.

We have demonstrated that two proteins previously predicted to be rhoptry kinases instead appear to be dense granule proteins. Importantly, both WNG1 and WNG2 contain the key kinase sequence motifs, suggesting they should be active within the dense granules and/or PV space. WNG1 (ROP35) was recently knocked out in Pru*Δku80* parasites, as part of a wider screen to identify the role of the predicted rhoptry kinases during chronic infection (44). Interestingly, although Δ*wng1* parasites retained the ability to differentiate into bradyzoites *in vitro*, parasites lacking this kinase formed approximately 75 % fewer tissue cysts within the brains of infected mice, suggesting WNG1 plays a role in the development or persistence of bradyzoites (44). Kinases have already been extensively explored as effector proteins in *Toxoplasma*, ranging from the secreted ROP16 and ROP18, to the recently described (dense granule protein kinases) annotated as ROP21 and ROP27 (45). While these kinases have been characterised at the host-parasite interface, no phosphatases had been reported at this location. In light of this, a novel finding in this study was the localisation of GRA44 at the PVM. This is the first example of a phosphatase that lies at the interface of this important boundary. Interestingly, GRA44 has a CRISPR score of −3.28 (42), lower than ASP5’s score of −1.45, which suggested that this protein is important for *in vitro* growth. This CRISPR score was validated as Δ*gra44* parasites exhibited a growth defect *in vitro*, a phenotype that has not previously been observed for ASP5-cleaved dense granule proteins, including GRA16 and MYR1, despite parasites lacking both of these proteins displaying a substantial virulence defect *in vivo* (14, 37). Finally, GRA44 was originally predicted to localise to the IMC based on an antibody that recognised the cytoplasmic face of the IMC, whose target was named IMC2 (46). The current version (v34) of ToxoDB does not recognise a protein termed IMC2, and the sequence used to immunise mice to generate the original IMC2 antibody is no longer annotated as part of IMC2A.

To assess the role of these newly described ASP5 substrates *in vivo*, we employed a mouse model to assess *Toxoplasma* infection in C57BL/6 mice. As a control, we inoculated mice with up to 50,000 PruΔ*ku80*Δ*asp5* (Δ*asp5*) tachyzoites and observed complete attenuation in virulence, while mice infected with proportional numbers of WT parasites succumbed during this time period. To validate the role of ASP5 cleavage in virulence for WNG1, WNG2 and GRA43, we infected mice with an RRL⟶ARL mutations, or knockouts. Surprisingly for each substrate we observed no significant difference in virulence for mice infected with WT parasites compared to RRL⟶ ARL parasites. These data suggest that at least for these non-exported proteins, efficient ASP5 processing is not required for their trafficking to the dense granules, PV and/or PVM. This however, does not discount that a small amount of protein may still be cleaved and this is enough to fulfil its function.

It is clear that there are more ASP5 substrates than we have identified. Not all ASP5 substrates were detected in our analyses, including GRA16 and MYR1. There are a several potential reasons for this, with two of the most likely being: 1. The ASP5-cleaved protein may be of low abundance in sampled parasites, and 2. The subsequent peptide difficult to enrich or not suitable for detection by mass spectrometry. The former point is supported by a recent study utilising TAILS to identify substrates of ASP3 in *Toxoplasma* (43). ASP3 processes microneme and rhoptry proteins that are generally more abundant than the dense granule proteins, potentially enabling greater substrate identification. Furthermore, it is likely that ASP3 substrates greatly outnumber ASP5 substrates. Overall, whilst it is clear that using two different methodologies (ie TAILS and HYTANE) to enrich for N-terminal peptides was beneficial (38, 39) it is clear that other methodologies will need to be adopted to identify a more ASP5 substrates.

In conclusion, we have used N-terminomics to identify novel substrates of the important protease ASP5, and validated a subset of these through tagging and mutagenesis of the endogenous locus within parasites. Deletion of several of these corresponded to a reduction in virulence during murine infection, but not to the same extent as infection with Δ*asp5* parasites, suggesting the dramatic reduction in virulence during Δ*asp5* infection results from many effectors that are not trafficked correctly. Overall, our data validate the essential role of ASP5 during infection and that the conserved RRL motif is critical for cleavage by this protease.

## Materials and Methods

### *Toxoplasma* transfection and growth

Candidate genes were tagged endogenously within parasites using the CRISPR/Cas9 system which has been adapted for use in *Toxoplasma* (47, 48). Briefly, genes were tagged just prior to the endogenous stop codon following guide selection from EuPaGDT (http://grna.ctegd.uga.edu/batch_tagging.html). The CRISPR target plasmid (made by Q5 mutagenesis, NEB) was co-transfected with homologous repair constructs containing, Ty- or HA-epitope tag as previously described (20). Two oligos with at least 30 bp of complementarity at their 3’ end, usually over the HA- or TY-epitope, were annealed together in IDT-duplex buffer by heating to 98 °C for two minutes then gradually allowed to cool (47). 10 ug of Cas9 plasmid was combined with the total 80 ug of annealed oligos, amd resuspended in 20 uL P3 solution (Lonza), in an Amaxa 4D Nucleofector (Lonza) using the code FI-115 (Human Unstimulated T-cells). Human foreskin fibroblasts were grown to confluency in Dulbecco’s Modified Eagle Medium (DMEM), supplemented with 10% Cosmic Calf serum (Hyclone) and refreshed with DMEM with 1% fetal calf serum when inoculated with parasites.

### PCR and plasmid construction

Please consult supplementary text for details.

### Immunofluorescence Assays (IFAs) and microscopy

Fixation, preparation of reagents and mounting for IFAs were performed as previously described (20). All images were captured on the Zeiss LSM 880 with Airyscan Detector. Before each session, channel alignment was performed using a FocalCheck fluorescence microscope test slide #1 (ThermoFisher) and subsequent images were automatically aligned in FIJI using the plugin TransformJ Translate.

### Western Blots

Immunoblot samples were pelleted then lysed for 30 mins at 4 °C in 1% v/v Triton-X 100 (brand), 1 mM MgCl_2_ in PBS (Gibco) supplemented with final 1 × complete protease inhibitors (Sigma) and 0.2 % v/v Benzonase (Merck). Samples were then combined with an equal volume of 2x Sample buffer and 20 μl loaded onto a gel. Proteins were transferred onto nitrocellulose (brand) then blocked in 5 % w/v milk in 0.05 % Tween 20-supplemented PBS. Primary and secondary antibodies were diluted in milk/PBS. For LI-COR Western blots, the membranes were imaged on an Odyssey Fc imager (LI-COR Biosciences) using IRDye 800CW goat α-rat, IRDye 800CW goat α-mouse and IRDye 680RD goat α-rabbit antibodies. Antibodies used in this study were: αHA 3F10 (Roche), αHA (Rabbit, in house), αTY1 BB2 (ThermoFisher), αGAP45 (49), αGRA1 (50), αMAF1 (13), αIMC1 (G. Ward, University of Vermont) and αROP2/3/4 (51).

### Quantitative Proteomics

Please consult supplementary text for details on SILAC labelling, protein extraction, TAILS and HYTANE purification, pH fractionation, LC MS/MS and data analysis

### Virulence Experiments

All mouse experiments were conducted within the regulations of the Walter and Eliza Hall’s Animal Ethics Committee (AEC). Pru*Δku80Δ*hx (WT), Pru*Δku80ΔhxΔasp5* (*Δasp5*), Pru*Δku80ΔhxΔgra43* (*Δgra43*), Pru*Δku80ΔhxΔwng1* (*Δwng1*) and Pru*Δku80ΔhxΔwng2* (*Δwng2*) parasites were grown in HFFs until ~10 % lysed, then scraped and released from host cells by passage through a 27g needle. Parasites were then counted and resuspended at the indicated dose in 200 μl PBS then injected intraperitoneally into 6 × 6-8-week-old C57BL/6 mice. Mice were weighed daily and culled after exhibiting bodyweight loss of greater than 15 % for three consecutive days if they were not going to recover (as per the WEHI AEC) or when moribund. All surviving mice were sacrificed 30 days after infection and the resulting cardiac bleed serum was used as a 1/50 dilution primary antibody to check for seroconversion against tachyzoite lysate.

### Datasets

The mass spectrometry proteomics data have been deposited to the ProteomeXchange (52) Consortium via the PRIDE (53) partner repository with the dataset identifier PXD008574.

Reviewer account details:

Username

Password: nHec97Mb

## Acknowledgements

We would like to thank Carolina Alvarado from WEHI Bioservices for her handling and maintenance of the infected mice. Ellen Yeh from Stanford for data and protocol sharing and for graciously accepting Michael Coffey into her laboratory where learnt the TAILS methodology with Katrina Hong. John Boothroyd, Mike Panas and Nicole Marino from Stanford for continual sharing of unpublished data, and the suggested nomenclature on the WNG kinases from Michael Reese at UT Southwestern.

